# A Method for Quantifying Molecular Interactions Using Stochastic Modelling and Super-Resolution Microscopy

**DOI:** 10.1101/177063

**Authors:** Keria Bermudez-Hernandez, Sarah Keegan, Donna R. Whelan, Dylan A. Reid, Jennifer Zagelbaum, Yandong Yin, Sisi Ma, Eli Rothenberg, David Fenyö

**Affiliations:** Dept. of Biochemistry and Molecular Pharmacology, New York University School of Medicine, New York, NY; Institute for Systems Genetics, New York University School of Medicine, New York, NY

## Abstract

We introduce the Interaction Factor (IF), a measure for quantifying the interaction of molecular clusters in super-resolution microscopy images. The IF is robust in the sense that it is independent of cluster density, and it only depends on the extent of the pair-wise interaction between different types of molecular clusters in the image. The IF for a single or a collection of images is estimated by first using stochastic modelling where the locations of clusters in the images are repeatedly randomized to estimate the distribution of the overlaps between the clusters in the absence of interaction (IF=0). Second, an analytical form of the relationship between IF and the overlap (which has the random overlap as its only parameter) is used to estimate the IF for the experimentally observed overlap. The advantage of IF compared to conventional methods to quantify interaction in microscopy images is that it is insensitive to changing cluster density and is an absolute measure of interaction, making the interpretation of experiments easier. We validate the IF method by using both simulated and experimental data and provide an ImageJ plugin for determining the IF of an image.

## Introduction

A fundamental question that many fluorescence microscopy experiments are trying to answer is whether or not the molecules under study interact and how this interaction changes under varying experimental conditions^1^. With the development of super-resolution (SR) fluorescence microscopy techniques, it is now possible to study biological samples at the sub-diffraction level, allowing researchers to observe molecules and their interactions at the tens of nanometers scale^2–6^. Since interactions are not directly observed, the spatial overlap, or co-localization between molecules is used as a surrogate for interaction. The co-localization observed at the scale provided by SR is more likely to represent true interaction, creating a need for improved methods of analysis. However, co-localization can occur at random and change with molecule density^7^. For example, differing experimental conditions can cause an increase in density and thus an increase in co-localization, while the interaction between molecules does not change. Here, we introduce a measure that takes into account this randomness and is not affected by changes in density.

In order to measure co-localization, there are generally two types of methods: intensity-based and object-based methods. Intensity-based methods focus on the correlation of pixel intensity levels in the color channels of the image^1, 8, 9^. These methods can be affected by noise^10^ and therefore rely on the correct subtraction of background pixel intensities. In addition, an increase of molecular density can cause an increase in the value of these measures^1^. They can also be difficult to assess for statistical significance when compared to randomized images since it is difficult to recreate the autocorrelation of pixels present in an experimental image^1, 7, 9, 11^.

Object-based methods identify objects in an image in order to quantify co-localization^8, 9, 12^. Image segmentation techniques^8, 13–17^ can be used to delineate objects or alternatively, each object can be represented by a point, such as the centroid. When the entire object is outlined, measures of object overlap can be used for analysis and randomized images produced for testing statistical significance^13, 14^. In addition, the degree of co-localization can be further quantified by comparing the distributions of overlap measurements from experimental data to that of simulations that model an increased probability of attraction^18, 19^. If objects are represented by points/coordinates in an image, spatial point process analysis tools such as the nearest-neighbor distance ^20, 21^, and the cross-correlation function can be utilized^22–24^. Statistical significance is then calculated by distinguishing values of these second order statistics from the null hypothesis that points are randomly distributed^9, 12, 25^. These methods can be directly applied to the results of single molecule localization microscopy (SMLM)^26–28^ a type of SR microscopy that localizes individual fluorophores and yields particle coordinate lists rather than intensity images as output. However, these coordinate-based methods internally apply radial averaging and therefore do not account for irregularly shaped objects ^23^. In addition, the application of these methods can be time consuming and require a high level of experience in statistical techniques and computer programming ^23^. Additional methods have been recently developed specifically for evaluating co-localization in SMLM data where a measure of co-localization is calculated for each coordinate (based on radial density) and then the aggregated results are evaluated graphically^29, 30^.

We have developed a co-localization measure called the Interaction Factor (IF), which is based upon measuring the amount of overlap between segmented objects, i.e. clusters of molecules, in an image. It is a probability estimate between 0 and 1, where 0 indicates that the co-localization observed is due to random occurrence and 1 indicates that all objects are co-localizing. This new measure addresses many of the drawbacks with other methods: it makes a comparison to realistic random images, it is insensitive to cluster density, it is easy to use and fast to calculate, and it provides an absolute rather than a relative measure of interaction. We provide both an ImageJ plugin and a python package that implement the IF calculation.

## Results

### Description of the algorithm

The input to the IF algorithm is a two-color fluorescence microscopy image with segmented objects corresponding to the molecular clusters in the image, and optionally a corresponding region of interest (ROI) (Fig. 1a(i)-(ii)). Firstly, a series of images are simulated by random placement of both sets of clusters within the ROI. Then, the frequency that each cluster of one color (the reference color) co-localizes (overlaps with at least one pixel) with any cluster of the other color is averaged over the total number of simulations. This estimates the probability of random co-localization for clusters of the reference color. (Figure 1a(iii)-(iv)). Finally, the Interaction Factor (IF) for the image is calculated from Equation 1, which describes the percentage of overlapping clusters of the reference color as a function of the IF. This equation can be derived from the algorithm described in Supplementary Figure 1, which provides a method for placing clusters in an image where the probability of overlap is greater than that of random placement. The input to this algorithm is the IF, which determines the increased probability of overlap (see Online Methods for derivation):

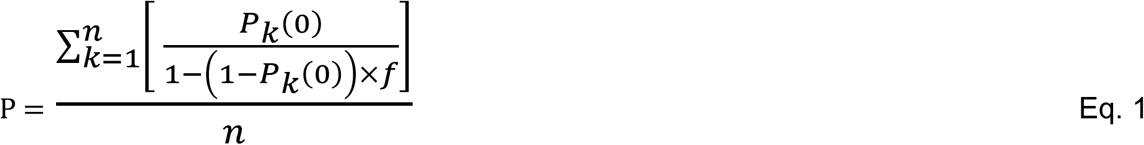

Where P = fraction of overlapping clusters of reference color, *f* = Interaction Factor (IF), *P*_*κ*_(0) = probability of overlap of the kth cluster in the randomized simulations and *n* = total number of clusters.

**Figure 1:**
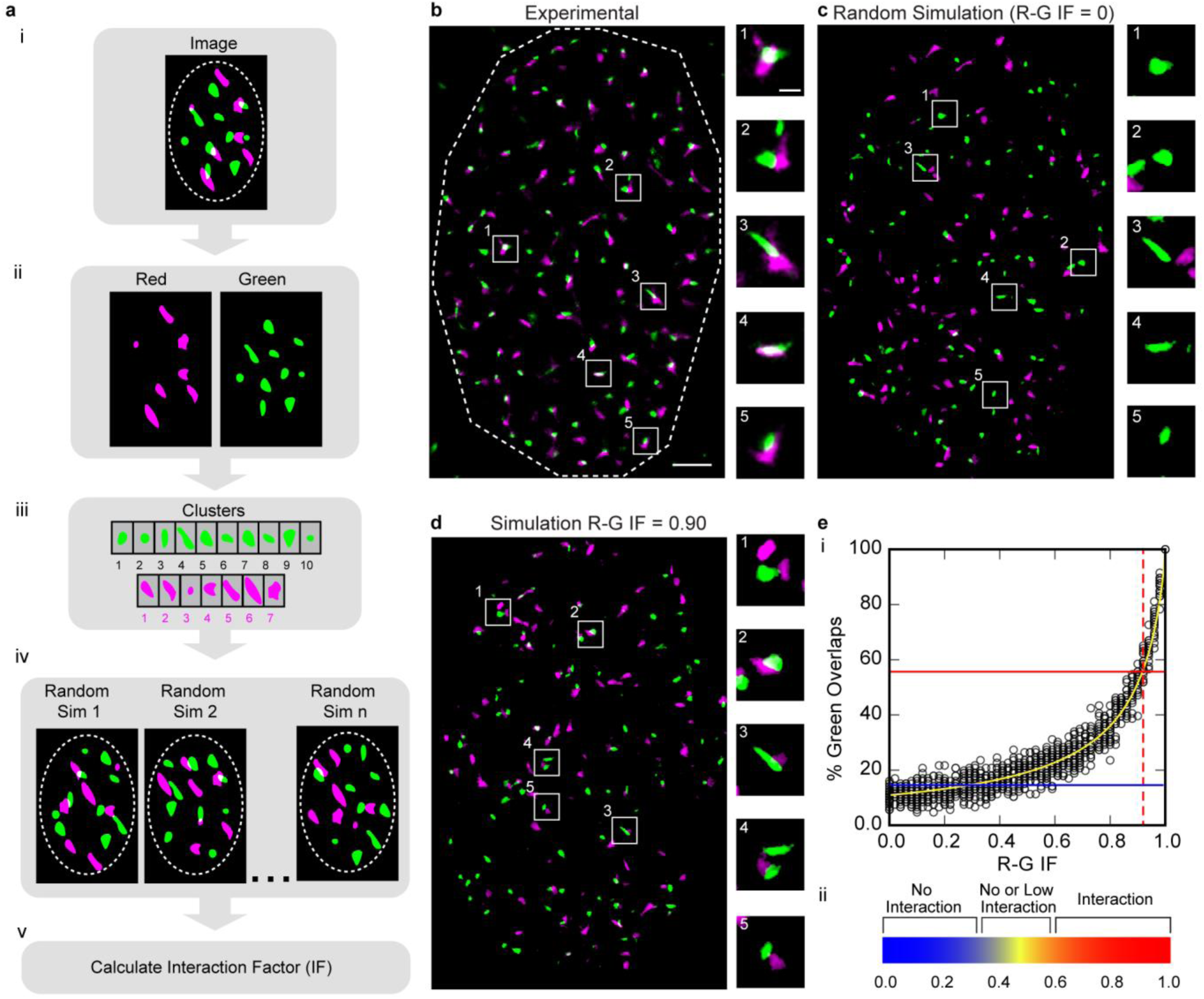
The Interaction Factor. (a) Schematic of the algorithm workflow. (a-e) For illustration purposes, the color red is represented as magenta. (a, i) Segmented two-color fluorescence microscopy (RGB) image of cell nucleus. ROI is outlined in dashed line. (a, ii) Color channels are separated. (a, iii) Clusters within the ROI are obtained from the segmented images. (a, iv) A simulation is created by placing clusters in random locations within the ROI. This process is repeated and the probability of green clusters (reference color) overlapping with red clusters is measured. (a, v) The probabilities are used to predict the IF (Equation 1). (b) Super resolution image (SMLM) of nucleus stained for two different proteins red and green. Dashed line indicates ROI, nucleus. Scale bar = 2µm. Boxes (1-5) point to clusters magnified in right column. Scale bar = 500 nm. (c) Simulation generated from clusters of experimental image (b) for IF = 0. Boxes point to same green clusters as experimental image (b) magnified in right column. Scale bar = 2 µm. (d) Simulation generated from clusters of experimental image (b) for Red-Green (R-G) IF = 0.90 (reference color = green). Boxes point to same green clusters as experimental image (b) magnified in right column. Scale bar = 2 µm. (e) The R-G IF curve. (e, i) Plot showing percentage of green cluster overlap for simulations generated with different R-G IFs. The yellow curve is a plot of its analytical prediction; points indicate values for individual simulations (n = 20 simulations per IF). The percentage of green overlap for the experimental image (b) was 56% (red line) resulting in a predicted R-G IF of 0.92 according to Equation 1. For a theoretical image with 15% green overlap (blue line), Equation 1 predicts R-G IF=0.22 (e, ii) Heat map that can be used as a guide to understand the meaning of the IF at different levels.

The IF can be calculated in the context of either color and therefore we specify the reference color used for the probability calculation, e.g. red-green (R-G) IF indicates that the probability is calculated using *green* as the reference color. The IF ranges from 0 to 1, where 0 indicates co-localization merely due to random occurrence and 1 indicates complete co-localization (all clusters co-localized). Figure 1b shows an SR image from experimental data of a cell nucleus with two fluorescently labeled proteins, red and green. Two scenarios with randomized (IF = 0, Fig. 1c) and highly correlated (IF = 0.9, Fig. 1d) placements of the two proteins are also shown, generated by the algorithm referenced above (Supplementary Fig.1 and Online Methods).

An alternative method to estimate the IF of the original image is to create groups of simulated images at IF levels ranging in steps from 0 to 1, and then find the IF that produces simulated images where the mean percentage of overlap of the reference color is closest to the percentage of overlap of the original image. For the experimental image in Figure 1(b), Figure 1e(i) shows the percentage of green co-localizing clusters for groups of simulated images (20 simulations per IF) plotted as a function of the R-G IF. The yellow curve is its analytical prediction (Eq. 1). The curve follows the data points of the simulations, confirming the derivation of Equation 1. When utilizing the IF to distinguish the degree of co-localization in an image from that of random occurrence, the diagram shown in Figure 1e(ii) can be used as a general guide.

### Validation and comparison to other methods

In order to determine if the R-G IF is affected by varying red cluster density, groups of simulated images with different numbers and sizes of red clusters were produced at R-G IF=0 (random), 0.25, 0.50, 0.75, 0.90 and 0.95 using the algorithmic procedure described in the Online Methods/Supplementary Fig. 1. The number of red and green clusters drawn in the simulated images (100) was taken from the average cluster counts of an experimental data set and the number of red clusters was then varied from 25 to 200. The size distributions of the clusters were based on measurements of segmented clusters from the experimental image shown in Figure 1b. These clusters were estimated as ellipses and drawn over the same ROI as shown for the experimental image and the sizes of red clusters were then varied by multiplying the axes of the ellipses by a factor of 0.5, 1.0, and 1.5 (while keeping the number of clusters the same). Both red cluster number and size were varied to cover a larger range of density levels than was observed experimentally.

The left panel of Figure 2a shows example simulated images with 25, 100, and 200 red clusters at IF=0 (green cluster number = 100, all groups). The right panel of Figure 2a shows example simulated images where the original cluster sizes were reduced by 50%, kept the same size (100%) and increased by 150% at R-G IF=0 (green cluster size unchanged and red/green cluster number = 100, all groups). Figure 2b(i) shows results of R-G IF calculations for images simulated at R-G IF=0 while varying red cluster number. In the left panel, both predicted R-G IF (left y-axis) and the percentage overlap of green clusters (right y-axis) are plotted. As the number of red clusters increases the percentage of overlapping green clusters increases (Kruskal-Wallis H test, all groups: p <0.0001), however there are no significant differences in calculated R-G IF for all groups (25 clusters: n=20, 0.14±0.17 (SD); 200 clusters: n=20, 0.04±0.08 (SD); One-way ANOVA, all groups: p=0.53). A similar result is shown for R-G IF=0.90 (25 clusters: n=20, 0.90±0.02 (SD); 200 clusters: n=20, 0.91±0.02; One-way ANOVA, all groups: p=0.25; Figure 2b(ii)). The right panels of Figure 2b(i) and (ii) show a more detailed view of each group of simulations. Results of varying the size of red clusters are shown in Figure 2b(iii) for R-G IF=0. Similar to the results of varying red cluster number, while the percentage of green overlapping clusters increases (One-way ANOVA, all groups: p < 0.0001), the calculated R-G IF accurately measures the simulated R-G IF for groups of images with different red cluster sizes, with no significant difference among groups (50% of size: n=20, 0.10±0.18 (SD); 150% of size: n=20, 0.10±0.13 (SD); One-way ANOVA, all groups: p=1.0). Simulations at R-G IF=0.90 also show that it is independent of red cluster size at this level (50% of size: n=20, 0.89±0.03 (SD); 150% of size: n=20, 0.90±0.02 (SD); Kruskal-Wallis H test, all groups: p=0.30; Figure 2b(iv)). These results illustrate that the IF is not affected by the density of clusters in the image.

**Figure 2:**
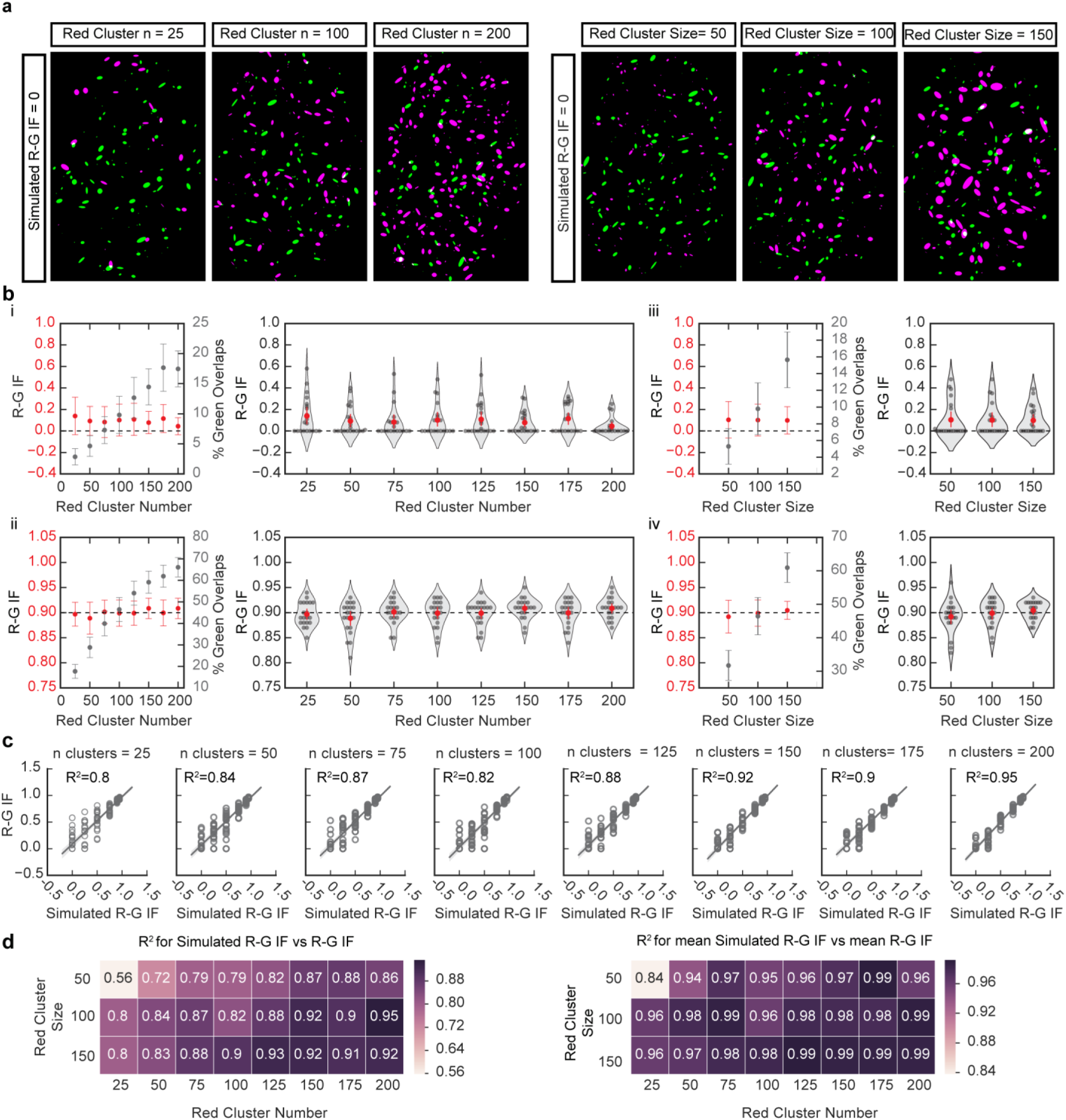
Effects of different cluster densities in R-G IF measurements and percentage of overlap. (a) Examples of simulations generated with 100 green clusters and different numbers of red clusters (left) or 100 red clusters of different sizes (right) for R-G IF = 0. For illustration purposes, the color red is represented as magenta. In generating the simulated images, cluster numbers and size distributions were sampled from experimental image (Fig. 1b). (b) Plots of percentage of green overlaps (gray) and calculated IF (red) for simulations generated with increasing red cluster number for R-G IF = 0 (i) and R-G IF = 0.90 (ii) show that the percentage of green overlaps increases with increasing red cluster number while the R-G IF remains constant. Plots of the calculated R-G IF for images generated with increasing red cluster sizes for R-G IF = 0 (iii) and R-G IF = 0.90 (iv) show that the percentage of green overlaps increases with increasing cluster size while the R-G IF remains constant. (b, i-iv) show that increasing red cluster number/size doesn’t affect the calculated R-G IF. (Means+/-SD, n = 20). (c) Plots of simulated R-G IF and calculated R-G IF for simulations with different red cluster number for five different R-G IFs. Coefficients of determination (*R*^*2*^) are shown as a measure of accuracy of the calculation (line, y=x; n = 20 images per R-G IF). (d) Heat maps of the *R*^*2*^ calculated from plotting the simulated R-G IF vs calculated R-G IF for simulations with different red cluster number and sizes show that the error range is greater in images with lower red cluster number and/or smaller size (left; n = 20 images per R-G IF). Heat maps of the *R*^*2*^ calculated from plotting the simulated R-G IF vs mean calculated R-G IF for the same simulations as left (right; n = 20 images per R-G IF).

Additional simulations were performed where green cluster density was varied (Supplementary Fig. 2) and both red and green cluster densities were varied simultaneously (Supplementary Fig. 3) and we found no dependence in the ranges tested between cluster density and IF. In addition to percentage of overlap, other common measures of co-localization in fluorescence microscopy, including the Manders’ coefficients^31^ and the Pearson’s coefficient also change with cluster density level (Supplementary Fig. 4).

Simulations at all R-G IF values measured from 0 to 0.95 are shown in Figure 2c, where the coefficient of determination (*R*^*2*^) of a simple linear regression model (calculated R-G IF (y) = simulated R-G IF (x)) is shown for images with varying red cluster numbers. The *R*^*2*^ provides a summative measure of the accuracy of the calculated R-G IF for all simulated R-G IF values at each increment of cluster number. Simulations were also performed while varying red cluster size at all R-G IF values. Figure 2d shows the effects of varying both size and number of red clusters on the accuracy of the R-G IF calculation. The *R*^*2*^ is shown for each combination of size and number, calculated using the measurements for all images in each group (left panel) and the mean of each group (right panel). Note the low accuracy of the R-G IF calculation in the upper left corner of the heat maps in Figure 2d (*R*^*2*^ = 0.56, left panel and *R*^*2*^ =0.84, right panel) for sparse images (low cluster density).

### Precision and statistical significance

Since the IF is based on the probabilities of cluster overlap from a set of random simulations, the IF value can vary with repeated runs of the calculation for a single image. To illustrate the degree of variation that can be expected, we generated simulations over a range of IFs and then repeated the IF calculation 20 times for each simulated image. The range of variation in the calculated IF for a random image was 0.10, while the range of variation in the calculated IF for an image at IF=0.96 was 0.01 (higher variation occurred in IFs closer to random), Supplementary Table 1.

While calculating the IF of a single image can be informative for preliminary work, it is recommended to draw conclusions based on a larger data set. In addition to data set size, the ability to distinguish between data sets is also dependent upon the IF value: the calculation is more precise for images with higher IFs. The IF curve (Fig. 1d) is relatively flat initially (near random levels of overlap), followed by a steeper section as the IF increases. Therefore, the IFs of images closer to random co-localization levels tend to have higher variance and are more difficult to distinguish: Figure 1e, blue line at a low green percentage of overlap (near random) intersects data points over a large range of x-values (simulated R-G IFs). Higher IFs have less variance and are easier to distinguish: Figure 1e, red line at a high green percentage of overlap intersects data points in a narrower range of x-values. Since the range of IFs that may give rise to a low level of overlap is larger, the IF for this type of data set can be better estimated by increasing the number of images measured.

In order to determine how the calculation of statistical significance is affected by the IF value for different data set sizes, we generated groups of images to simulate experimental data sets at a range of IF values where n=5, 10 or 20 images per group. The R-G IF was calculated for each simulated image and a Student’s t-test was performed between groups of images to obtain the associated p-values. To reduce the potential of bias from using a single experimental image to make the simulations, the comparison was repeated using identified clusters from 20 different images from an experimental data set, and the average p-value was calculated. The results of this comparison are shown in Fig. 3. The data sets that showed no significant difference are colored dark purple; note the larger proportion of this color on the heat map for smaller sized data sets, but also note the extent of the dark purple for groups with low R-G IF as opposed to groups with high R-G IF (left side of heat maps vs. right side of heat maps). Figure 3 can be used as a general guide for the number of images required for significance when gathering data, based on the expected or preliminary IF level.

**Figure 3:**
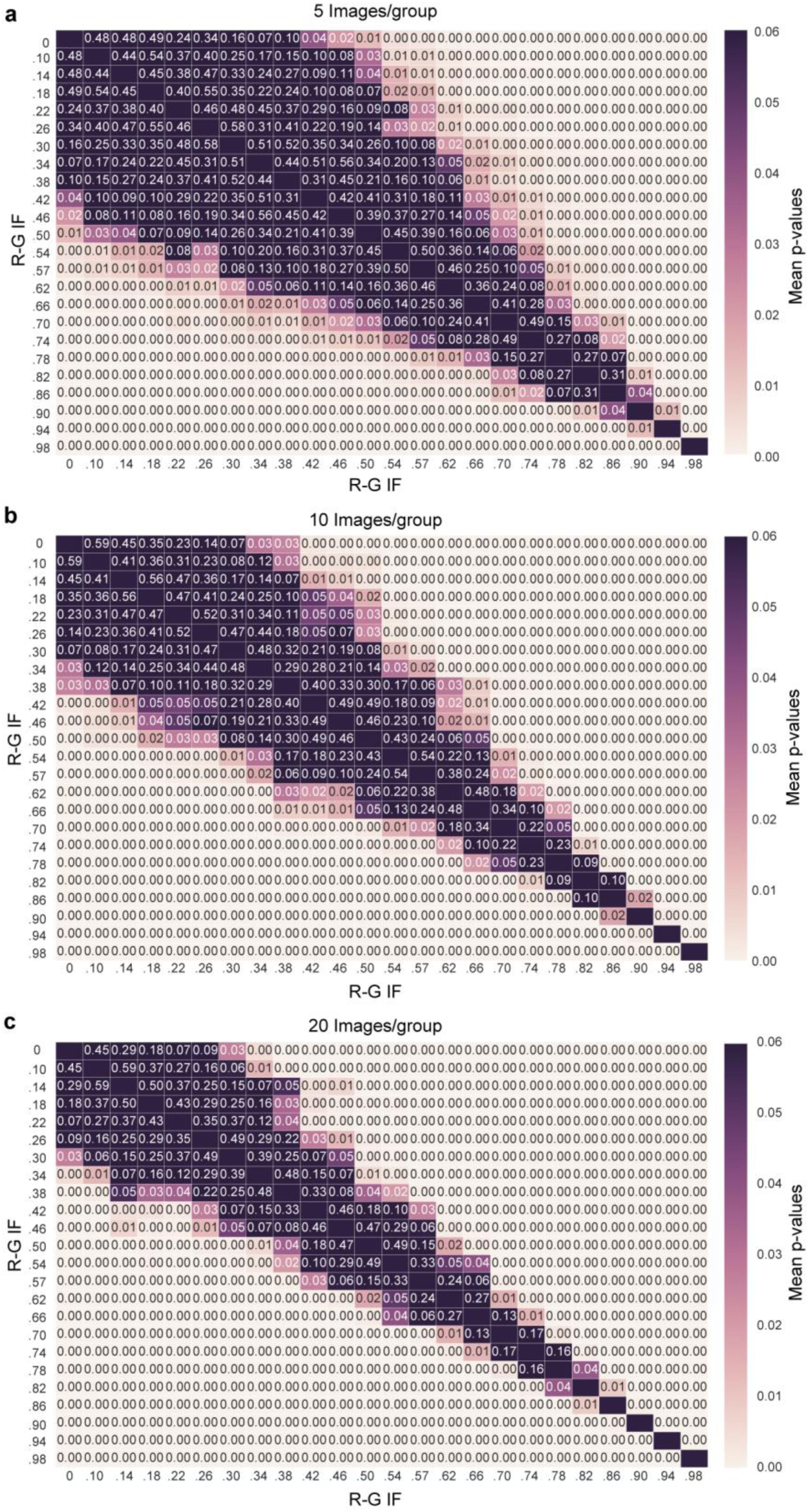
Statistical significance of R-G IF with different data sizes. (a) A heat map showing the mean Student’s t-test p-value that resulted from the comparison of the calculated R-G IFs of simulated experimental groups with 5 images/group for a range of R-G IFs (values indicate the mean p-value for 20 comparisons). (b) A heat map showing the mean Student’s t-test p-value that resulted from the comparison of the calculated R-G IFs of simulated experimental groups with 10 images/group for a range of IFs (values indicate the mean p-value for 20 comparisons). (c) A heat map showing the mean Student’s t-test p-value that resulted from the comparison of the calculated R-G IFs of simulated experimental groups with 20 images/group for a range of R-G IFs (values indicate the mean p-value for 20 comparisons).

### Analysis of the interaction of DNA damage repair proteins

Following the development of SR microscopy, researchers have successfully devised assays to use SMLM to probe the spatial and temporal recruitment of DNA damage repair proteins to double stranded breaks (DSBs) ^32^. We believe the IF approach could bring new insights to the protein-protein interactions involved in key aspects of both non-homologous end joining (NHEJ) and homologous recombination (HR), the principal DSB repair pathways^33^. We tested the IF approach in three experimental datasets that looked at damaged and undamaged cells and included proteins believed to be involved in HR or NHEJ^33^. For datasets 1 and 2 (Fig. 4a, left panel), U2OS cells were synchronized post-mitosis, released into G1 and treated with 2.5 ng/mL neocarzinostatin (NCS) (DNA-damaged) or normal media (control) for 30 minutes, cells were fixed and immunolabeled for phosphorylated DNA-PKcs (pDNA-PKcs) and DNA Ligase IV (LigIV) (Fig. 4b) or phosphorylated ATM kinase (pATM) and LigIV (Fig. 4c). For experimental dataset 3 (Fig. 4a, right panel), U2OS cells were synchronized post-mitosis, released for 26 hours to establish a predominantly G2-phase cell population and treated with NCS (DNA-damaged) or normal media (control) for 30 minutes. After 30 minutes of recovery, cells were fixed and immunolabeled for BRCA1 and BRIP1 (Fig. 4d).

**Figure 4:**
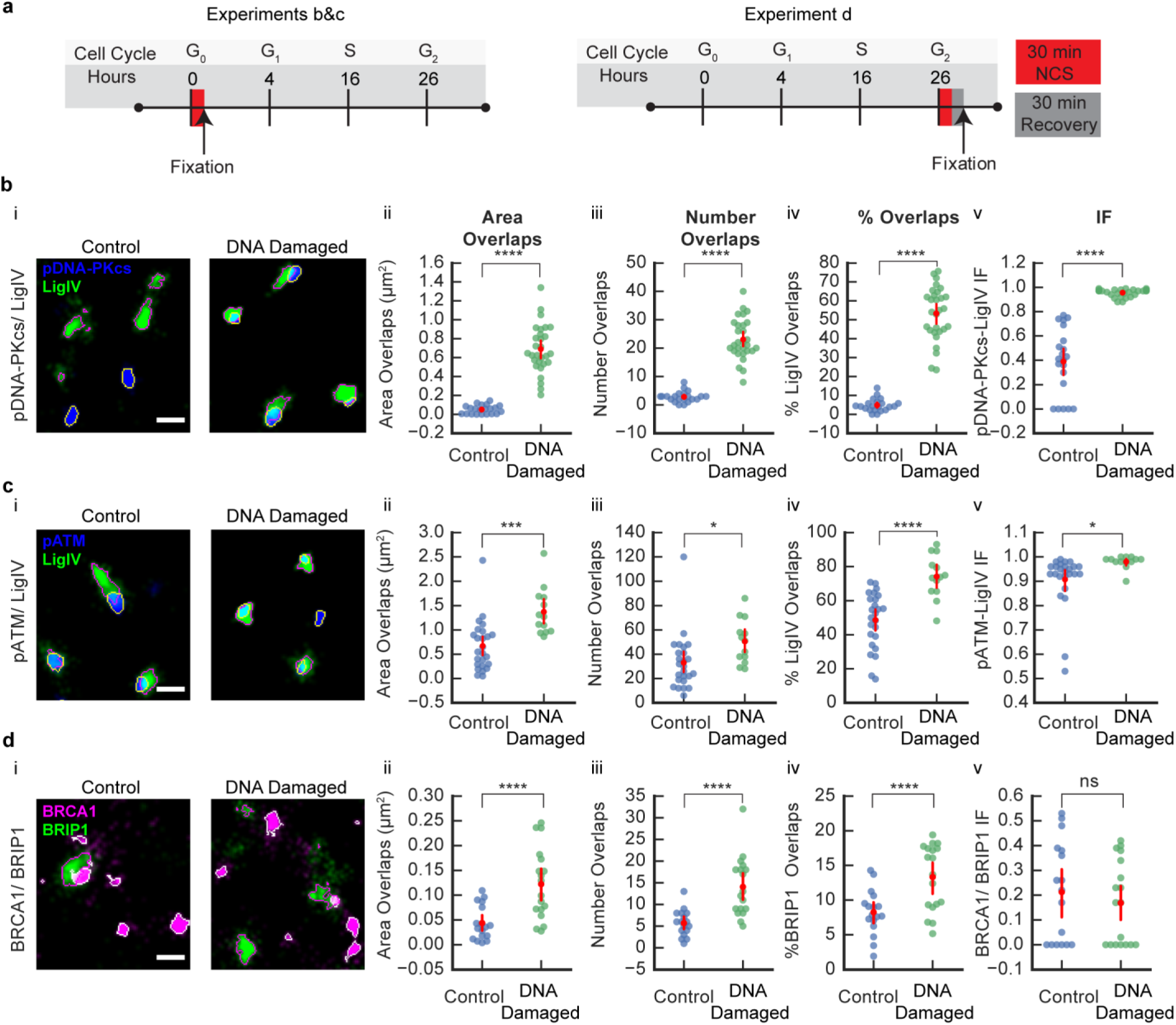
Comparison of the R-G IF for SMLM images of U2OS cell nuclei labeled for proteins involved in NHEJ and HR. (a) A schematic of the experimental timeline for NCS treatment and fixation of U2OS cells quantified in part b&c (left) and part d (right). (b, i) pDNA-PKcs and LigIV. Examples of an area of SMLM images of U2OS cells in control (left) and after NCS treatment (DNA-damaged; right) labeled for LigIV (green; outlined magenta: segmented clusters) and pDNA-PKcs (blue; outlined yellow: segmented clusters). Scale bar = 400 nm. (b, ii) Area of overlaps between pDNA-PKcs/LigIV was greater in DNA-damaged compared to control group. (b, iii) Number of overlaps between pDNA-PKcs/LigIV was greater in DNA-damaged compared to control group. (b, iv) Percentage of LigIV clusters overlapping with pDNA-PKcs was greater in DNA-damaged compared to control group. (b, v) IF between pDNA-PKcs/LigIV was greater in DNA-damaged compared to control group. Groups: control (n = 21); DNA-damaged (n = 29). (c, i) pATM and LigIV. Examples of an area of SMLM images of U2OS cells in control (left) and after NCS treatment (DNA-damaged; right) labeled for LigIV (green; outlined magenta: segmented clusters) and pATM (blue; outlined yellow: segmented clusters). Scale bar = 400 nm. (c, ii) Area of overlaps between pATM/LigIV was greater in DNA-damaged compared to control group. (c, iii) Number of overlaps between pATM/LigIV was greater in DNA-damaged compared to control group. (c, iv) Percentage of LigIV clusters overlapping with pATM was greater in DNA-damaged compared to control group. (c, v) IF between pATM/LigIV was greater in DNA-damaged compared to control group. Groups: control (n = 25); DNA-damaged (n = 13). (d, i) BRCA1 and BRIP1. Examples of an area of SMLM images of U2OS cells in control (left) and after NCS treatment (DNA-damaged; right) labeled for BRIP1 (green; outlined magenta: segmented clusters) and BRCA1 (magenta; outlined white: segmented clusters). Scale bar = 400 nm. (d, ii) Area of overlaps between BRCA1/BRIP1 was greater in DNA-damaged compared to control group. (d, iii) Number of overlaps between BRCA1/BRIP1 was greater in DNA-damaged compared control group. (d, iv) Percentage of BRIP1 clusters overlapping with BRCA1 was greater in DNA-damaged compared to control group. (d, v) IF between BRCA1/BRIP1 was not significantly different after DNA-damage compared to control group (n.s. = not significant; p>0.05). Groups: control (n = 17); DNA-damaged (n = 18). (b-d) Error bars represent 95 CI. *p<0.05, ***p<0.001 ****p<0.0001 by t-test.

#### pDNA-PKcs and LigIV

For experimental dataset 1, a Welch’s t-test showed that the area of overlap between pDNA-PKcs and LigIV was significantly greater after DNA-damage compared to the control group (control: n=21, 0.05±0.01*µm*^*2*^(SEM); DNA-damaged: n=29, 0.69±0.05*µm*^*2*^(SEM); Welch’s t-test, p<0.0001; Fig. 4b, ii). Similarly, the number of overlaps between pDNA-PKcs and LigIV clusters was significantly greater after DNA-damage compared to control group (controls: 2.86±0.41(SEM); DNA-damaged: 23.03±1.34(SEM); Welch’s t-test, p < 0.0001; Fig. 4b, iii). To determine if the proportion of LigIV clusters that overlapped with pDNA-PKcs changed after NCS treatment, we calculated the percentage of clusters that overlapped with pDNA-PKcs and found a significant increase in the DNA-damaged group (control: 4.85±0.73(SEM); DNA-damaged: 53.30±2.62(SEM); Welch’s t-test, p < 0.0001; Fig. 4b, iv).

These data could be interpreted as both proteins being closer in space caused by an increase in protein-protein interaction. It is also possible that the increase in these measurements is simply due to an increase in cluster area or number. The total area covered by LigIV and pDNA-PKcs was greater after DNA-damage compared to control group (Supplementary Fig. 5a) raising the possibility that the observed changes in overlap measurements were caused by an increase in cluster area. The effect of changing cluster number is inconclusive given that while the number of pDNA-PKcs clusters increased, the number of LigIV clusters decreased after DNA-damage (Supplementary Fig. 5a).

We calculated the IF based upon LigIV as the reference protein (pDNA-PKcs-LIgIV IF) and found that the mean IF was greater after DNA-damage compared to control group (control: 0.39±0.06; DNA-damaged: 0.96±0.01(SEM); Welch’s t-test, p<0.0001; Fig. 4b, v), suggesting an increase in interaction between LigIV and pDNA-PKcs after DNA damage. The mean IF of 0.39±0.06 (SEM) suggests that without NCS induced DNA damage there is no or very little interaction between LigIV and pDNA-PKcs. Whereas, the mean IF for the DNA-damaged group of 0.96±0.01(SEM) suggests that there is very high interaction between LigIV and pDNA-PKcs after DNA damage. Furthermore, we found that the choice of reference protein (LigIV or pDNA-PKcs) had no effect on the results (Supplementary Fig. 6a).

#### pATM and LigIV

For experimental dataset 2, a Student’s t-test showed that the area of overlaps between pATM and LigIV was significantly greater after DNA-damage compared to control group (control: n = 25, 0.67±0.10*µm*^*2*^(SEM); DNA-damaged: n=13, 1.37±0.14*µm*^*2*^(SEM); Student’s t-test, p<0.0001; Fig. 4c, ii). Similarly, the number of overlaps between pATM and LigIV clusters was significantly greater after DNA-damage compared to control group (controls: 33.04±4.56(SEM); DNA-damaged: 50.54±5.01(SEM); Student’s t-test, p=0.02; Fig. 4c, iii). To determine if the proportion of LigIV clusters that overlapped with pATM changed after NCS treatment, we calculated the percentage of LigIV clusters that overlapped with pATM and found a significant increase in the DNA-damaged group (control: 48.67±3.26(SEM); DNA-damaged: 74.26±3.51(SEM); Student’s t-test, p<0.0001; Fig. 4c, iv). Altogether these data suggest that there is an increase in the interaction between LigIV and pATM, however it is also possible that the increase in overlap measurements is due to an increase in the number or area of clusters. Additional data (Supplementary Fig. 5b) showed no differences between control and DNA-damaged groups in cluster area or number, which would suggest that the observed increase in overlap is due to an increase in protein-protein interaction.

We calculated the IF based upon LigIV as the reference protein (pATM-LigIV IF) and found that the mean IF was greater after DNA-damage compared to control group (control: 0.91±0.02(SEM); DNA-damaged: 0.98±0.01(SEM); Welch’s t-test, p=0.03; Fig. 4c, v). Further examination of the IF measurements show that the mean IF was 0.91±0.02(SEM) for the control group suggesting that without the NCS DNA damage induction there is high interaction between LigIV and pATM. Whereas, the mean IF for the DNA-damaged group was 0.98±0.01(SEM) suggesting that the interaction was already present in the absence of DNA damage induction, and increased slightly after DNA damage. As in the previous data set, we found that the choice of reference protein (LigIV or pATM) had no effect on the results (Supplementary Fig. 6b).

#### BRCA1 and BRIP1

For experimental dataset 3, a Student’s t-test showed that the area of overlaps between BRCA1 and BRIP1 was significantly greater after DNA-damage compared to control group (control: n=17, 0.04±0.10*µm*^*2*^(SEM); DNA-damaged: n=18, 0.12±0.02*µm*^2^(SEM); Welch’s t-test, p<0.0001; Fig. 4d, ii). Similarly, the number of overlaps between BRCA1 and BRIP1 clusters was significantly greater after DNA-damage compared to control group (controls: 5.71±0.73(SEM); DNA-damaged: 14.06±1.61(SEM); Welch’s t-test, p<0.0001; Fig. 4d, iii). To determine if the proportion of BRIP1 clusters that overlap with BRCA1 changed after NCS treatment, we calculated the percentage of BRIP1 clusters that overlapped with BRCA1 and found a significant increase in the DNA-damaged group (control: 8.26±0.78(SEM); DNA-damaged: 13.33±1.12(SEM); Student’s t-test, p<0.0001; Fig. 4d, iv). Altogether, these data suggest that there is an increase in protein-protein interaction or that the respective clusters increased in number and/or area. Additional data (Supplementary Fig. 5c) showed the DNA-damaged group exhibited an increase in both BRIP1 and BRCA1 cluster area and number raising the possibility that this increase in cluster density caused the observed increase in overlap.

We calculated the IF based upon BRIP1 as the reference protein (BRCA1-BRIP1 IF) and found no significant difference in IF between control and DNA-damaged group (control: 0.21±0.05(SEM); DNA-damaged: 0.17±0.04(SEM); Student’s t-test, p=0.49; Fig. 4d, v), further supporting the hypothesis that the increase in cluster density caused the increase in overlaps. Moreover, the low mean IFs in both groups would suggest that the proteins are not interacting before or after DNA induced damage during G2 phase. As in the previous data set, the choice of reference protein (BRIP1 or BRCA1) had no effect on the results (Supplementary Fig. 6c).

## Discussion

The Interaction Factor (IF) is calculated by using stochastic simulations to determine the probability for random overlap and deriving the relationship between IF, the experimentally observed overlap and the probability for random overlap (Eq. 1). Using both simulated and experimental data we show that the IF is widely applicable to a large range of situations encountered in super-resolution microscopy.

In order to further elucidate the meaning of the IF in a biological context, we compared it to the equilibrium dissociation constant (K_d_) for receptor-ligand binding, based on the assumption that the observed co-localization in an image is the result of binding between the two molecules. We calculated *in silico* K_d_ values for simulated images with increasing numbers of clusters of one color (corresponding to increasing the concentration of the ligand), and then fit a saturation binding model to the results to obtain K_d_ values and compared these with calculated IF values. As expected, there is an inverse relationship between K_d_ and IF (Supplementary Fig. 7). When K_d_ decreases, the probability for molecules to be in a bound state increases, leading to increased co-localization and increased IF.

The inherent variability in the amount of cluster overlap when there is no interaction or only weak interaction causes significant variation in the IF measurement for low IFs (Figure 1e(i), Supplementary Table 1). While increasing the size of the data set can mitigate this problem somewhat (Figure 3), we recommend that in general reporting interaction for IF levels below 0.5 is done with caution (Figure 1e(ii)). In the case where the calculated IF is reported as 0, this implies that the level of co-localization is equal to or less than that found for the random case. We do not attempt to distinguish the IF for the case where the co-localization is less than that found for random in our current model, as we have yet to encounter this situation biologically.

For our simulated data, the accuracy of the IF calculation was lowest when using 25 clusters for both colors, especially when the clusters were small in size. Therefore, we do not recommend using the IF calculation when images contain less than 25 clusters and they are small with respect to the size of the ROI. In addition, it is important that both sets of clusters are distributed uniformly over the ROI. If this is not the case, for example if one of the clusters of molecules is bound to a membrane, then the procedure can be modified by allowing one set of clusters to be left in their original positions while randomly placing only the second set of clusters. An extension that could be added to the IF algorithm would be to substitute the uniform distribution for randomization of clusters within the ROI with an arbitrary shape for the distribution of clusters.

In the simulations produced for the IF calculation, clusters were randomly placed in the ROI but we did not take the additional step of randomly rotating the clusters. While this is straightforward to add to the algorithm, we found that it would not significantly affect the results and therefore was an unnecessary complication (Supplementary Fig. 8).

The IF has the potential to probe the asymmetry of the interaction, i.e. for two interacting clusters of molecules, the IF for one set of clusters with respect to the other can be directional. For example, in the case where only a small subset of one protein interacts with the second protein while the second protein almost always interacts with the first, the IF will be different depending on the choice of the reference protein. This fact is taken into account in the formulation of the IF (Equation 1, Supplementary Fig. 1) and it is recommended to calculate the IF in the context of both sets of clusters to fully describe the meaning of the co-localization in the image. In our ImageJ plugin (Supplementary Fig. 9), both calculations are always reported.

The method for determining co-localization was chosen as the percentage of overlapping clusters where overlap is defined as any amount of pixel overlap between segmented objects. This seemed a valid basis for measuring co-localization in SR images, but we recognize that the optimal measure can be different depending on the application. Our model may be adjusted to consider co-localization in a different manner, for example a particular amount of pixel overlap, the overlap of a cluster centroid with another cluster or a minimum distance of clusters.

The current implementation of IF is for interactions in two dimensions, but it would be straightforward to extend it to three dimensional images by simple implementation changes without any need to change the basic definition of the IF.

Finally, we recognize that the algorithm is not suited for images where molecules cannot be arranged in clusters, and thus other co-localization methods such as cross-correlation analysis ^22, 23^, the CBC method^29^ or intensity-based methods ^1, 8, 9^ are better suited for this type of data.

In summary, we have introduced a new technique to evaluate co-localization measurements in microscopy images, the Interaction Factor (IF). It provides an absolute measure of the interaction between proteins and because it is insensitive to cluster density, it can directly be used for calculating the effective size of changes between experimental conditions. The absolute nature of the IF makes interpretation of co-localization in microscopy images easier by allowing biologists not only to quantify the size of the change in interaction but also to distinguish the case where the change goes from no interaction to high interaction from the case where the interaction is already high but then increases even further. The examples we have shown here are all from dSTORM images but the method is equally applicable to other SR and conventional fluorescence microscopies, or for that matter, to any images with overlapping objects of different types.

## Methods

### Sample preparation and Immunofluorescence

For experimental data sets 1 and 2 (Fig. 4a, left panel and Fig. 4b&c), U2OS (American Type Culture Collection) cells were cultured routinely in McCoy’s 5A medium supplemented with 10% fetal bovine serum and penicillin/streptomycin. Cells were synchronized post-mitosis by serum starvation for 72 hours, before release into G1 and treatment with 5 ng/mL of neocarzinostatin (NCS) for 30 minutes (DNA-damaged) or normal media (control). For experimental dataset 3 (Figure 4a, right panel and Fig. 4d), U2OS cells were synchronized by serum starvation for 72h, released for 16 hours and treated with CPT for 30 minutes (DNA-damaged) or normal media (control).

Cells were then subsequently prepared for imaging via cytoplasmic extraction with cold CSK buffer containing 0.5% Triton X-100, washed with DPBS (with calcium and magnesium), and fixed for 15 minutes in 4% paraformaldaheyde in DPBS. Coverslips were blocked with blocking solution [20 mg/mL BSA, 0.2% gelatin, 2% (wt/vol) glycine, 50 mM NH4Cl, and PBS], then stained with primary antibodies dilutions specified by the manufacturer either overnight at 4 °C or for 1 h at room temperature) and secondary antibodies (usually at 1:1,000–5,000 for 30 min at room temperature) in blocking solution before imaging. The following primary antibodies were used: phosphorylated DNA-dependent protein kinase catalytic subunit (pDNA-PKcs; ab18356, Abcam), DNA Ligase IV (LigIV; ab26039, Abcam) and phosphorylated Ataxia telangiectasia-mutated protein (pATM; ab36810, Abcam), Breast cancer type 1 susceptibility protein (BRCA1; sc6954, Santa Cruz) and Fanconi anemia group J protein (BRIP1; NB100-416, Novus). Anti-mouse and anti-rabbit secondary antibodies conjugated to Alexa Fluor 647, 568 and 488 were used for visualization (Thermofisher).

### Microscopy Setup for Single-Molecule Imaging

We used a custom-built microscopy setup based on a Leica DMI3000 microscope equipped with an HCX PL APO 63× NA 1.47 OIL CORR TIRF objective, followed by achromatic 2× tube lens magnification. The microscope was coupled to 473 nm (200 mW), 532 nm (200 mW), and 640 nm (150 mW) diode-pumped solid-state lasers to excite the sample in a Highly Inclined and Laminated Optical Sheet (HILO) illumination mode for improved signal to noise ratio and to reject out of plane fluorescence. Sample emission was collected and split into multiple channels through the use of proper dichroic and emission narrow-band bandpass filters in conjunction with the use of a Photometrics DV2 multichannel imaging system to image two colors simultaneously, side-by-side, onto a single EM-CCD camera (Andor iXon+ 897) acquiring at 33 Hz.

For accurate alignment and mapping of the two color channels, we first imaged diffraction-limited fluorescent beads that have wide emission spectra spanning all channels (Life Technologies). The location of the beads was matched for each channel, and a mapping matrix was generated using a custom mapping routine (IDL; Exelis Visual Information Solutions). In brief, this routine is based on the use of a polynomial morph-type mapping function, in which mapping coefficients are generated by Gaussian and centroid fits to the subdiffraction limit point spread functions of the fluorescence beads.

### SR Imaging

SR imaging was achieved through a modified direct stochastic optical reconstruction microscopy (*d*STORM) approach^18^. Images were acquired in HILO illumination mode^34^. Using these acquisition modes and the filters described earlier, we found only negligible fluorescence cross-talk between channels. Subdiffraction two-color colocalization error was <20 nm in our instrument^18, 35^. SR imaging conditions were achieved through the addition of 100 mM MEA and an additional oxygen-scavenging system (1 mg/mL glucose oxidase, 0.02 mg/mL catalase, and 10% glucose)^28, 36^. Movies containing a minimum of 2,000 frames were used to generate reconstructed super-resolved images.

### Analysis

SR image reconstruction was performed using the freely available QuickPALM plug-in for ImageJ^37^. We specifically used an FWHM of 4 and a signal-to-noise ratio (SNR) of 2-4 to detect localized particles from our raw data. Localized molecules were written to rendered images by the QuickPALM plug-in with a pixel size of 20 nm. For the experimental datasets, ROIs were manually drawn to outline cell nuclei using ImageJ. Then, images were segmented with the Otsu thresholding method and all clusters less than 4 pixels in area were excluded from the analysis.

### Derivation of IF formula

Suppose we have an image with sets of clusters from two different color channels. Clusters of one color are first placed randomly on the image (random color). Then, clusters of the second color (reference color) are placed according to the procedure outlined in Supplementary Figure 1 and Online methods (see below). Based on the steps in the figure, a formula can be derived to relate the IF to the percentage of overlapping clusters (of the reference color), with the only additional parameter being the probability of overlap for each cluster in a completely random simulated image (IF=0). The table below illustrates the probability of overlap for cluster k (1 ≤ k ≤ n), (reference color) at the 1^st^ and 2^nd^ iteration of the algorithm, where *P*_*κ*_(0) = Probability of overlap of cluster k where both sets of clusters placed randomly and f = Interaction Factor.

*For cluster k:*

| Iteration (i) | Probability of Overlap | Probability of No Overlap | Probability of Continuing |
| --- | --- | --- | --- |
| 1 | $P_k(0)$ | $1 - P_k(0)$ | $(1 - P_k(0)) \times f$ |
| 2 | $P_k(0) \times [(1 - P_k(0)) \times f]$ | $(1 - P_k(0)) \times [(1 - P_k(0)) \times f]$ | $(1 - P_k(0)) \times [(1 - P_k(0)) \times f] \times f$ |

From the above table, we can derive *P*_*κ*_(*f*), the probability of overlap of cluster k at IF = f, and then simplify:

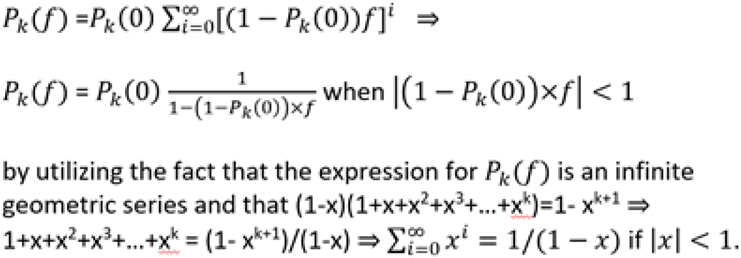

Finally, where P = % of overlapping clusters (reference color) in the experimental image, we have:

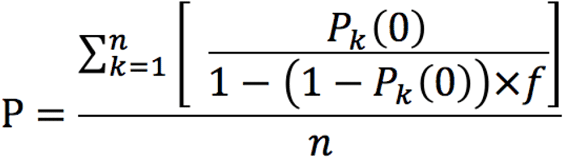

### Generation of IF simulations

To generate IF simulations, clusters of one color are placed randomly on a new image within the boundaries of the ROI, if present. Then, clusters of the second color (reference color) are placed according to the flowchart (Supplementary Fig.1). This process is repeated for each cluster: (1) the cluster is placed randomly within the ROI. (2) If the IF is equal to 0, then the cluster is kept in that position. (3) If the IF is not equal to 0, the program determines if the cluster is overlapping with one or more pixels of a cluster of the other color. (4) If it is overlapping, then the cluster is kept in that position. (5) If instead it’s not overlapping, then a random number between 0 and 1 is generated. (6) If that number is greater than the IF, the cluster is kept in that position. (7) Conversely, if the random number is not greater than the IF, then the process is repeated from step (1) above. The random number generation utilized in the python package is random.randint() from the python standard library. The random number generation utilized in the ImageJ plugin was Math.random() from the Java standard library.

### Statistics for Ellipse Simulations and Experimental data comparisons

Ellipse simulations data is expressed as mean ± SD. Experimental data is expressed as mean ± SEM. The p criterion was 0.05. Tests for equality of variances were determined by performing a Bartlett’s test. For analyses of two groups, if results showed significant differences in variances, Welch’s t-tests were performed. For analyses of more than two groups, if results showed significant differences in variances, Kruskal-Wallis H tests were performed. Linear regression, two-tailed Student’s t-tests, Welch’s t-tests, Kruskal-Wallis H tests, and One-way ANOVAs were performed using the Scipy Stats package^38^.

### Additional Measurements

In the text the **percentage of overlaps** refers to the percentage of clusters of the reference color overlapping with the other color. They were counted by first generating masks for both channels such that *C*_*mask*_ = *C > th*, were *C* is the color channel and *th* is the threshold value, followed by counting the clusters that co-localized with a cluster of the other color, and dividing it by the total number of clusters multiplied by 100. The **number of overlaps** were counted by first generating an overlap mask

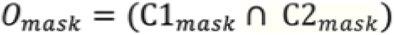

where *C1*_*mask*_ is the channel 1 mask, *C2*_*mask*_ is the channel 2 mask, followed by counting the number of overlapping regions in the overlap mask. The **area of overlaps** was calculated by adding the area of all the overlapping regions and multiplying it by the scale. Mander’s coefficient 1 (**M1**) was calculated by utilizing the following formula:

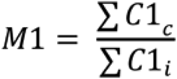

where *C1*_*c*_ is the intensity value of a pixel in channel 1 above threshold that co-localized with channel 2 above threshold and *C1*_*i*_ is the intensity value for a pixel in channel 1 above threshold. Mander’s coefficient 2 (**M2**) was calculated by utilizing the following formula:

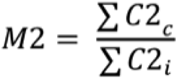

where *C2*_*c*_ is the intensity value of a pixel in channel 2 above threshold that co-localized with channel 1 above threshold. The **Pearson’s coefficient** was calculated using the following formula:

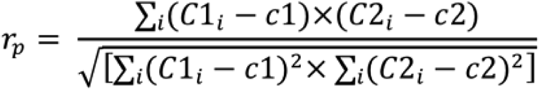

where *C1*_*i*_ is the intensity value for a pixel in channel 1, *c1* is the mean intensity value for channel 1, *C2*_*i*_ is the intensity value for a pixel in channel 2, and *c2* is the mean intensity value for channel 2.

### ImageJ Plugin and python package

We provide both an ImageJ plugin (Supplementary Fig. 8) and also a python package to calculate the IF for an image. The ImageJ plugin is called “Interaction Factor Package” and contains two dialog boxes. The first, called “Interaction Factor”, calculates the Interaction Factor for an image. The user can outline the ROI and choose the color channels to be examined before running the IF calculation. Built-in ImageJ thresholding methods may be applied to the image to obtain masks for the channels. The plugin allows the user to test different thresholding methods interactively by drawing an overlay based on the chosen method. Once satisfied, the user can then run the IF calculation. The user may choose to randomly arrange both sets of clusters or keep one set of clusters in their original positions while rearranging only the second set. The output is the calculated IFs for the image (in the context of both color channels, e.g. red-green IF and green-red IF), along with a “p-value” indicating the proportion of the random simulations produced in the IF calculation which had a percentage of overlap ≥ that of the original image. The number of random simulations performed by default in the plug-in is 50, providing a precision of 0.02 for the probability calculation. The second dialog box, called “Interaction Factor Simulations”, allows the user to create a set of simulated images at a chosen IF based on an input image. The interface for pre-processing the input image is similar to the “Interaction Factor” dialog box, except that the user may also choose the IF level and number of simulated images generated. The plugin source code and .jar file, as well as a detailed user guide are included as Supplementary Software. The python package “IFSimPy” provides analogous tools for calculating the IF as well as creating simulated images at any IF. The python source code is also included as Supplementary Software.

## Acknowledgements

K.B. was supported by the Molecular Oncology and Immunology training grant T32 CA009161. We would like to thank Mario Delmar and Esperanza Agullo-Pascual for the opportunity to collaborate and for comments on the ImageJ plugin. We would also like to thank Bart Aromando for his comments and work on the user guide for the plugin.

### Author Contributions

K.B., S.K., S.M. and D.F. conceived the method. D.R.W. provided invaluable input and feedback on the utility of the method and the functionality of the ImageJ plugin. D.R.W., D.A.R., J.Z., Y.Y. and E.R performed and/or supervised experiments. K.B. and S.K. performed the analysis. The manuscript was co-written by K.B., S.K. and D.F., with contributions from all authors.

### Competing Financial Interests

None.

